# Acidic monosaccharides become incorporated into calcite single crystals

**DOI:** 10.1101/2020.08.03.234310

**Authors:** Arad Lang, Sylwia Mijowska, Iryna Polishchuk, Simona Fermani, Giuseppe Falini, Alexander Katsman, Frédéric Marin, Boaz Pokroy

**Author notes:** A. Lang and S. Mijowska contributed equally to this work as first authors.

## Abstract

Carbohydrates, along with proteins and peptides, are known to represent a major class of biomacromolecules involved in calcium carbonate biomineralization. However, in spite of multiple physical or biochemical characterizations, the explicit role of saccharide macromolecules (long chains of carbohydrate molecules) is not yet understood in mineral deposition. In the present study we investigated the influence of two common acidic monosaccharides (MSs), which are the simplest form of carbohydrates and are represented here by glucuronic and galacturonic acids, on the formation of calcite crystals *in vitro*. We show that the size, morphology and microstructure of calcite crystals are altered when they are grown in the presence of these MSs. More importantly, MSs were found to become incorporated into the calcite crystalline lattice and induce anisotropic lattice distortions, a widely studied phenomenon in other biomolecules related to CaCO_3_ biomineralization but never before reported in the case of single MSs. Changes in the calcite lattice induced by MS incorporation were precisely determined by the technique of high-resolution synchrotron powder X-ray diffraction. We believe that the results of this research may deepen our understanding of the interaction of saccharide polymers with an inorganic host and shed light on the implications of carbohydrates for biomineralization processes.

## INTRODUCTION

Biocalcification, when considered as a process, can be viewed as the ability of living systems to synthesize calcified structures whose physical properties (morphology, size, structure, hardness, resistance to fracture, and optical properties) are tailored for one or more biological functions.^[1–6]^ This tailoring is possible only because of the complex interplay at the nanoscale level between inorganic prenucleation clusters of calcium carbonate and organic macromolecules that collectively form the skeletal matrix. Such interactions are indeed a sine qua non condition for driving biominerals along nonconventional crystallization pathways and for generating final products that differ significantly from their abiotically produced counterpart.^[7–9]^

Since proteins provide a direct link to the genetic machinery involved in calcification, a great deal of attention has been directed to skeletal matrix proteins and their interactions with calcium and/or calcium carbonate. Dozens of papers on the biochemical properties of skeletal proteins have been published over the last two decades,^[10,11]^ and complete skeletal protein repertoires are nowadays known in corals,^[12]^ brachiopods,^[13]^ molluscs^[14]^ and sea urchins.^[15]^ Many of these protein mixtures have been tested in vitro for their ability to inhibit the precipitation of CaCO_3_^[16]^ or to modify the size and morphology of calcite crystals.^[17]^ More surprisingly, proteins,^[18–20]^ or their most elementary building blocks, amino acids,^[21–24]^ were found to induce anisotropic lattice distortions in their calcitic, aragonitic or vateritic crystalline hosts. Several synthetic assays also showed that other molecules can get incorporated into calcite such as gels,^[25,26]^ diblock copolymer particles^[27]^ and many more.

Based on numerous studies, polysaccharides (PSs) are also recognized as major components of the organic matrices associated with biomineral formation, and they play a vital role in maintaining the structural integrity of the organisms. But this fraction, although quantitatively important, has unfortunately been neglected in biomineralization studies. This is largely because the primary structure of PSs is not encoded in the genome and, furthermore, their chemistry is even more complex and subtle than that of proteins: the several existing monomers may combine in linear or in branched polymers and, for a given polymer, the general rule is polydispersity of molecular weights, of charges, and of monomer compositions. The discovery of PSs in calcified tissues supplied clear evidence of two classes with strikingly different properties: chitin and charged PSs. The first one, representing an insoluble polymer of N-acetyl glucosamine, was found early in history to be a constituent of some mollusk shells,^[28]^ a fact that was confirmed much later.^[29–32]^ It consists of an insoluble organic matrix which forms a scaffold for the emerging mineral.^[33,34]^ Specific orientations of chitin fibers can promote the templating of CaCO_3_ in the carapaces and shells of marine animals.^[35,36]^ Negatively charged PSs, often referred to as “mucopolysaccharides”, consist of soluble macromolecules that were first detected by histological staining,^[37,38]^ and later by bulk biochemical characterization.^[39– 41]^ While chitin acts as an organic framework on which biominerals grow, mucopolysaccharides are thought to have several other functions, including interaction with growing crystals of calcium carbonate during the process of biomineralization. They also have been shown to control the formation and stabilization of the amorphous phase of CaCO_3_.^[42]^ Their occurrence in mollusk shell matrices led Addadi and coworkers^[43]^ to propose a molecular model where mucopolysaccharides, by concentrating calcium ions at the vicinity of the protein template, cooperate with acidic proteins. Later, the ability to obtain mild hydrolysis and the subsequent quantification of monosaccharides (MSs) by High-Performance Anion-Exchange chromatography using the Pulsed Amperometric technique allowed bulk characterization of monosaccharide contents of the skeletal tissues of cephalopods,^[44]^ bivalves,^[45]^ brachiopods,^[46]^ crustaceans,^[47]^ sea urchins^[48,49]^ and corals,^[50]^ which in some cases showed true monosaccharidic signatures. However, accurate characterization of PSs in terms of primary structure is still in its infancy in biomineralization studies, if we refer to the paucity of oligosaccharide sequences obtained so far.^[51,52]^

The purpose of this study was to evaluate the key role played by saccharides in biocalcification processes and to show that they are effective in inducing lattice distortions in calcite crystals. To this end we focused on two acidic MSs, glucuronic acid (GlcA) and galacturonic acid (GalA). These two chemical species are the uronic acids that are most commonly found in biological systems. They derive from their respective precursors, glucose and galactose, via oxidation of the 6^th^ carbon atom (of the carbon ring) to carboxylic acid, thereby creating the so-called uronic acid pathway.^[53]^ Under physiological pH conditions, with pK values of 3.2/3.3 (GlcA) and 3.51 (GalA),^[54]^ they are deprotonated, meaning that they carry a net negative charge and are consequently prone to interact with inorganic cations. The calcium-binding abilities of GlcA and GalA have been investigated in detail,^[55]^ as have those of their homopolymers, the poly-GlcA and poly-GalA acids,^[56]^ which tend to form gels in the presence of divalent cations. GlcA is abundantly present in connective tissues, where it is one of the two hexose residues that constitute the elementary motif of the chondroitin sulfate polymer^[57]^ or of hyaluronic acid,^[58]^ both found in cartilage and bone tissues. GalA is less abundant than GlcA in connective tissues but is particularly enriched in pectins, the complex sugar family found in plant cell walls. GlcA has been detected in the skeletal matrix of several calcifying metazoans including corals,^[50,59]^ brachiopods,^[46]^ two crayfishes *(Pacifastacus leniusculus* and *Orconectes limosus*),^[60]^ the land snail *Helix aspersa maxima*,^[61]^ the freshwater snail *Lymnaea stagnalis*,^[62]^ the cephalopod *Nautilus macromphalus*^[63]^ and also Nautilin-63,^[64]^ and the sea urchin *Arbacia lixula*.^[48,49]^ In the same samples the occurrence of GalA is irregular, since it has been detected only in the shell matrix of *N. macromphalus*^[63]^ but not in that of Nautilin-63^[64]^, in one crayfish only (*O. limosus*)^[60]^ and in the test matrix of the sea urchin *A. lixula*.^[48,49]^

In non-metazoan biocalcifications, uronic acids have been well described as monomeric constituents of PSs of coccoliths (the calcified plates of coccolithophore algae). In particular, a PS enriched in D-galacturonic acid was identified in *Emiliania huxleyi*^[65]^ and its primary structure was solved.^[66]^ Further biochemical characterization showed that the galacturonic acid residues, via their carboxyl groups―and not via the ester sulphate groups―are solely responsible for the inhibition of CaCO_3_ precipitation,^[67]^ a property that establishes uronic acids as key players in the interaction with calcium carbonate. Later, the presence of uronic acids in the PSs of coccoliths was detected in several strains and species.^[68–70]^ Interestingly, two recent papers attribute lattice distortions in calcite to the incorporation of coccolith macromolecules.^[71,72]^ However, none of the reports firmly establishes the types of the molecules―purified or not―responsible for this effect. To fully understand such polysaccharide−mineral interactions we need to start at the molecular level, as we did for proteins.

In this paper we describe the growth of calcite crystals under both ambient and hydrothermal (HT) conditions and in the presence of MSs, bearing in mind that the incorporation of simple carbohydrates into the crystal lattice of calcite has not been previously reported. There is no doubt that elucidation of this hitherto unfathomable mechanism could provide conclusive evidence and clarifications with regard to specific interactions between PSs and a crystalline host. We show here that GlcA and GalA, as monomers, can become incorporated in relatively large amounts, and accordingly, might contribute substantially to the lattice distortions in biogenic calcite.

## RESULTS AND DISCUSSION

To determine the feasibility of MS incorporation into the calcite lattice we utilized the vapor diffusion (VD) method and grew calcite crystals in the presence of 0.15 M of GlcA or GalA under ambient conditions (see Experimental Section). Calcite crystals were collected after 48 h and characterized. High-resolution scanning electron microscopy (HRSEM) clearly revealed changes in the morphologies and shapes of calcite crystals grown in the presence of each of the MSs compared to the traditional rhombohedral shape of the pure calcite (Figure 1). The sizes and morphologies of the crystals (see low magnification images in Figure S1) are representative of the whole crystal population. Calcite-GalA crystals retained the rhombohedral structure, but displayed rough and deformed facets (Figure 1b). Interestingly, calcite-GlcA crystals exhibited the elongated scalenohedral-like morphology (Figure 1c) resembling that observed for calcite crystals containing high amounts of incorporated amino acids, in particular aspartic acid.^[23]^ As also seen in Figure 1, calcite-GalA crystals reached a size of about 1 mm, whereas calcite-GlcA crystals were about 100 μm in size. These first observations indirectly indicated that MSs probably become incorporated into the lattice of calcite crystals.

**Figure 1.**
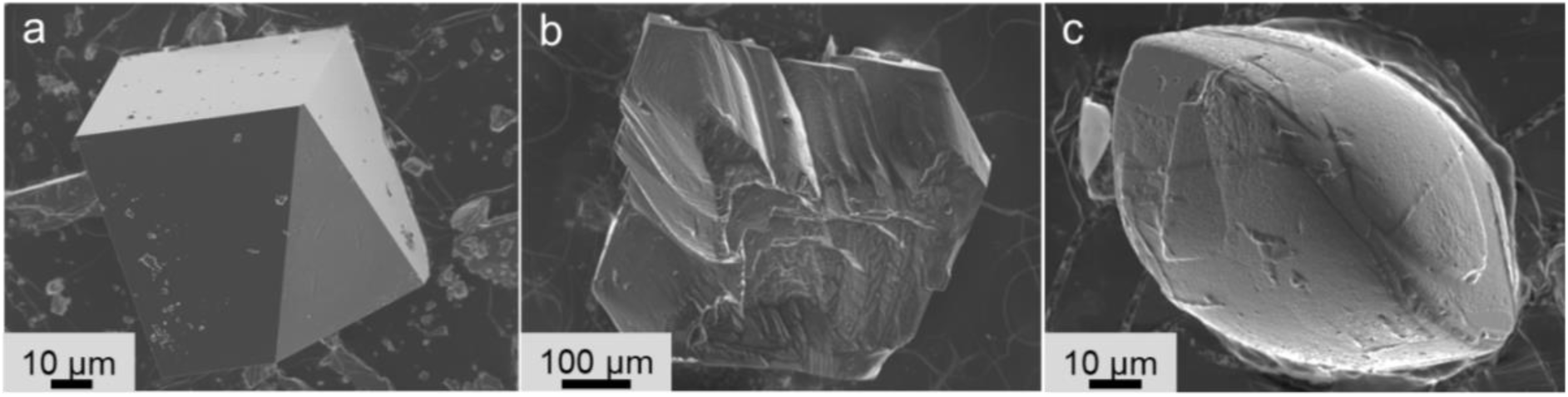
HRSEM shows calcite crystals prepared by the vapor diffusion (VD) method. (a) A pure calcite crystal. (b) In the presence of GalA and (c) of GlcA.

Our recent study of the growth of calcite crystals in the presence of amino acids and under hydrothermal (HT) conditions indicated that both high temperature and high pressure promote the incorporation of amino acids and lead to significantly higher levels of incorporation than those achieved under ambient conditions.^[23]^ We, therefore, attempted to further increase the loading of MSs, by utilizing the HT method (see Experimental Section). Calcite growth experiments were performed at 134 °C and water pressure of ∼0.3 MPa for 4 h. To follow the concentration-dependent effects, we added GlcA or GalA at concentrations of 0.075, 0.1 or 0.15 M to the reactant solution. As can be seen from Figure 2, changes in the morphology of calcite crystals grown via HT synthesis were indeed governed by the concentration of the added MSs. Calcite crystals precipitated at lower GlcA and GalA molarities (0.075 and 0.1 M) maintained the rhombohedral morphology, though the crystal facets displayed rough facets (Figure 1a, b, d, e). At the highest concentration (0.15 M) crystal morphologies were akin to that of scalenohedral calcite with elongated facets (Figure 2c, f), similar to that observed in the case of aspartic incorporation^[23]^ via the HT method.

**Figure 2.**
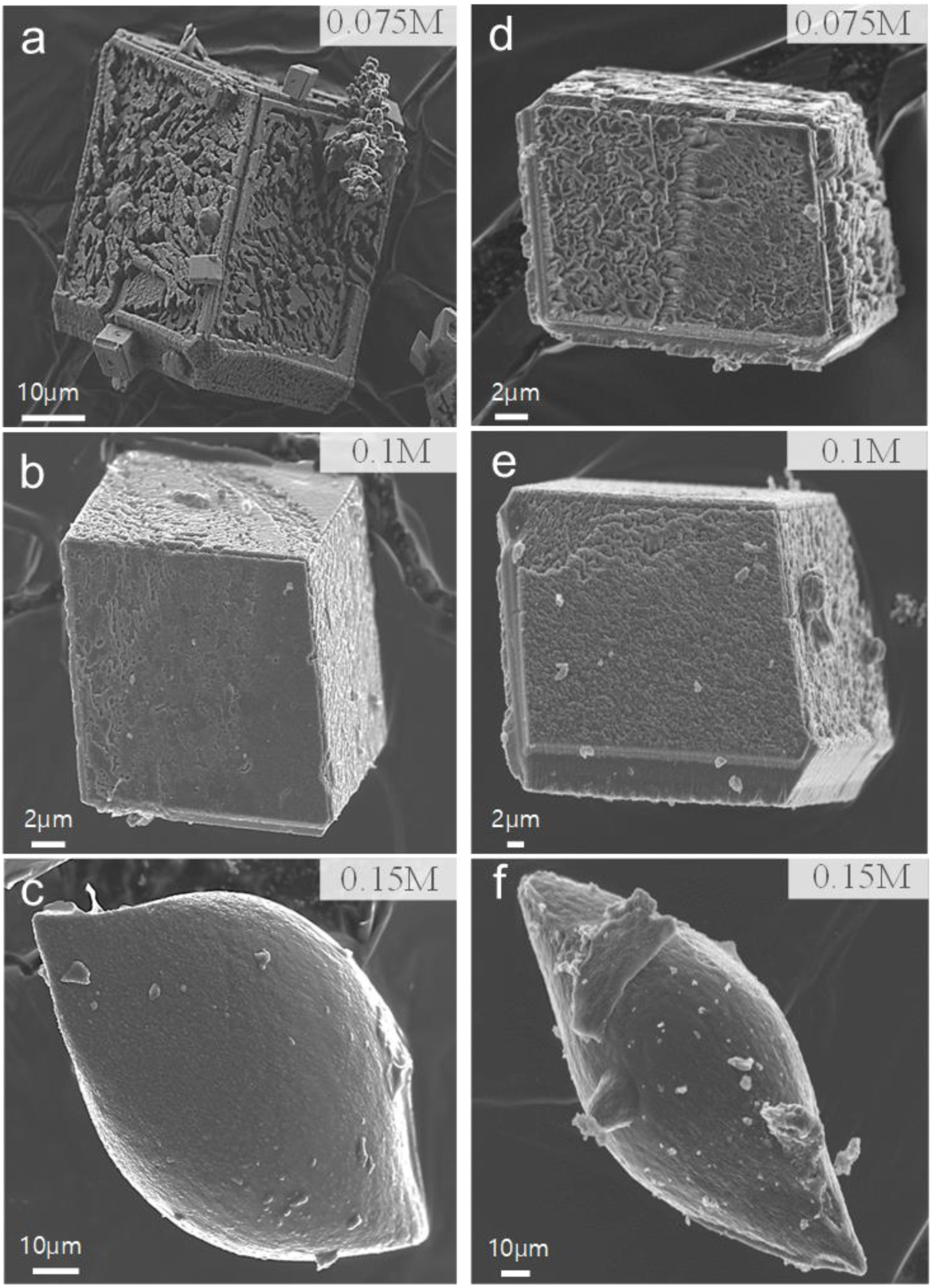
HRSEM shows calcite crystals synthesized by the hydrothermal (HT) method in the presence of different concentrations of MSs. (a−c) Calcite with 0.075, 0.1 and 0.15 M of GalA, respectively. (d−f) Calcite with 0.075, 0.1 and 0.15 M of GlcA, respectively.

We employed synchrotron high-resolution powder X-ray diffraction (HRPXRD) to precisely determine the lattice parameters of crystals grown in the presence of GlcA or GalA via both VD and HT methods. This technique has been shown to be the most sensitive and accurate method to determine the presence of intracrystalline molecules.^[18,20,21,73]^ Diffraction patterns were collected from as-grown samples and from samples annealed ex-situ at 300 °C for 2 h. Initial data analysis showed that in the case of both growing methods the samples containing MSs were pure calcite, and that only control samples consisted of calcite together with small amounts of vaterite impurity (Figure S2). The (104), (006) and (110) diffraction peaks were explicitly analyzed to evaluate the influence of MSs on the *c*- and *a*-parameters of the calcite unit cell. Figure 2 summarizes the evolution of (104), (006) and (110) diffraction peaks, both before and after annealing, of the calcite samples grown via HT or VD in the presence of 0.15 M of GlcA or GalA. In the case of both growing methods, our first observation was the presence of shifts in the positions of all diffraction peaks relative to those of the calcite control (Figure 2, Figure S4). Moreover, the higher the concentration of MSs in the reactant solution, the larger the shift in positions of the diffraction peaks (see Figure S3 for the case of the HT method). These observations strongly supported our suggestion that MSs indeed become incorporated into the crystalline structure of calcite and induce distortions in its lattice. However, as observed by the magnitude of the diffraction peak shifts, the level of incorporation is higher in the case of the HT than of the VD method of synthesis, a phenomenon we have already reported in the case of aspartic acid incorporation.^[23]^ Moreover, while (104) and (006) diffraction peaks demonstrate a shift towards lower 2*θ* upon incorporation of MSs via the VD and HT methods (Figure 3a,b and Figure S3, S4), in the case of calcite crystals grown via the HT method solely, the (110) diffraction peak shifts in the opposite direction and towards higher 2*θ* angles (Figure 2c,d). These findings imply that whereas the calcite lattice expands in all crystallographic directions when incorporation occurs under ambient conditions (i.e. VP method), under HT growth conditions, a shrinkage occurs along the a-axis only. Similar to our observation in the case of aspartic acid incorporation, this lattice distortion pattern stems from the fact that when the concentration of incorporated MSs is high, a large expansion occurs along the *c-*axis thereby contracting the *a*-axis.

**Figure 3.**
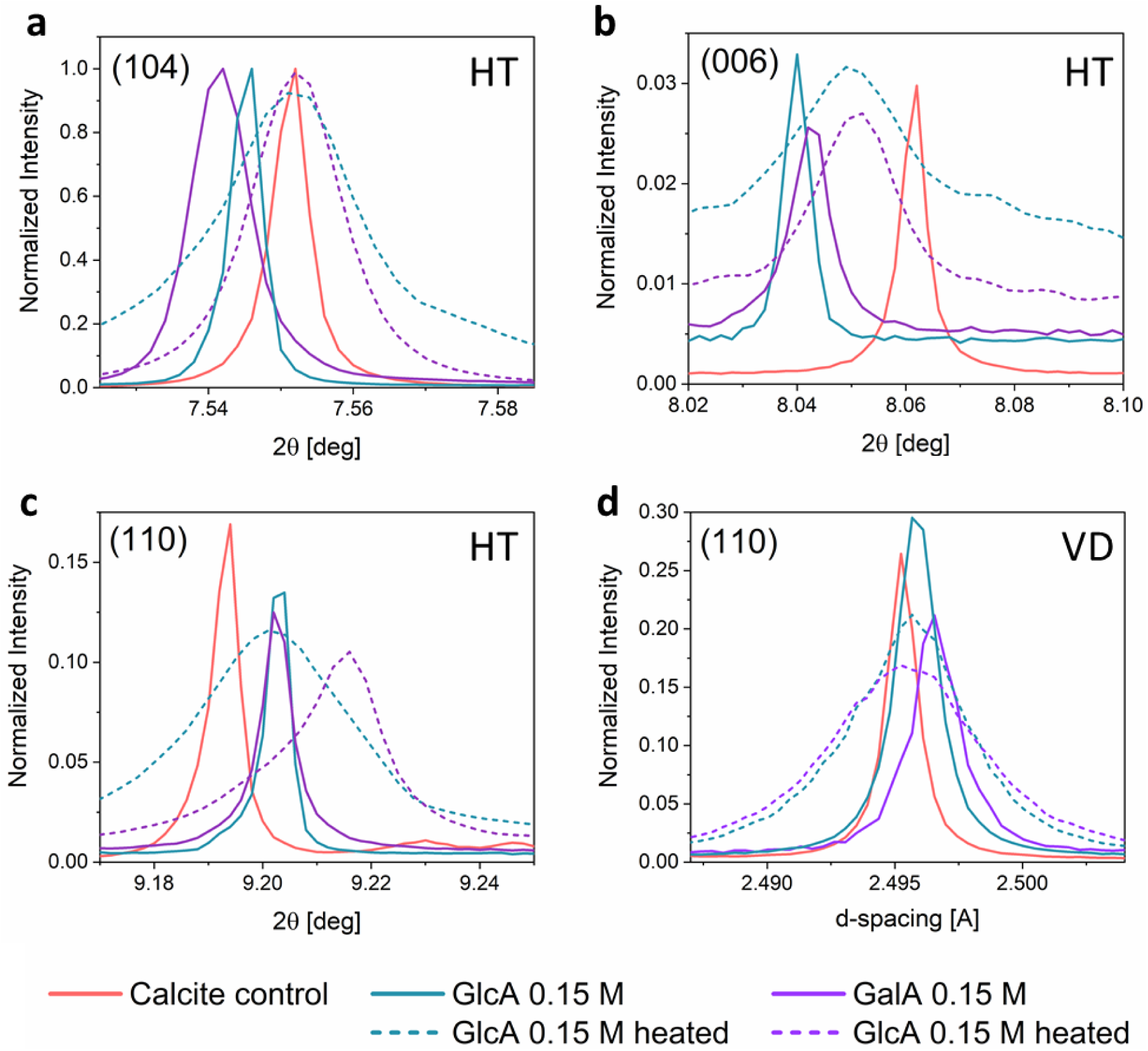
Evolution of the (a) (104), (b) (006) and (c) (110) diffraction profiles in the case of calcite crystals grown via the HT method, and (d) (110) diffraction profile in the case of calcite crystals grown via the VD method. GalA or GlcA concentration in the reactant solution is 0.15 M in the case of both syntheses. XRD data were collected from as-grown crystals (solid lines) and after their annealing (dashed lines) at 300 °C for 2 h and at a wavelength of 0.3999 Å.

As a comparison we grew calcite in the presence of 0.15M glucose. This MS molecule as opposed to GalA and GlcA MSs does not contain any acidic carboxylic acid function. Interestingly in this case negligible lattice distortions were detected (see Figure S5 and table S1). This emphasizes the importance of the acidic group in facilitating incorporation of MSs into the lattice of calcite probably via the carboxylate group substituting the carbonate group^[74]^ or interacting with calcium ions.

Annealing of the samples caused significant broadening of the diffraction peaks and relaxation of the lattice distortions, as shown by the shift in diffraction peaks towards the position of the control calcite sample (Figure 3). Even though such temperature-induced relaxation is a common feature in crystals that incorporate organic molecules, the lattice distortions caused by incorporated MSs under HT conditions were not fully removed here after annealing. This can be seen from the (006) and (110) diffraction peaks before and after annealing (Figure 3b, c). It is evident that MSs are more stable with temperature than amino acids, which readily decompose to CO_2_ and NH_3_ upon heating.^[75]^ In the case of MSs the carboxylic acid probably decomposes first, as amino acids, whereas the sugar molecule, being much more stable,^[76]^ remains intact within the calcite lattice and does not allow for full lattice relaxation. Rietveld refinements performed on the collected XRD data enabled us to determine the lattice parameters of all the studied crystals (Table S1). Lattice distortions were calculated utilizing the equations and, where *m* is MS-incorporated calcite, is the reference (pure) calcite, and *j* = 0.075 M, 0.1 M, or 0.15 M (see ref. ^[18]^). Figure 4 provides quantitative results of the calculated lattice distortions induced upon incorporation of GlcA or GalA via both synthesis methods. The magnitude of lattice distortions along the *c*-axis increases with the concentration of MSs in the solution, reaching 0.278% and 0.249% for 0.15 M GlcA and 0.15 M GalA, respectively. In fact, these lattice distortion values along the *c*-axis are larger than those reported for biogenic calcite.^[18]^ The *a*-lattice parameter slightly expands with low MS concentrations (0.075 and 0.1 M), but shows significant contractions of −0.104% and −0.103%, respectively, upon addition of 0.15 M of GlcA or GalA.

**Figure 4.**
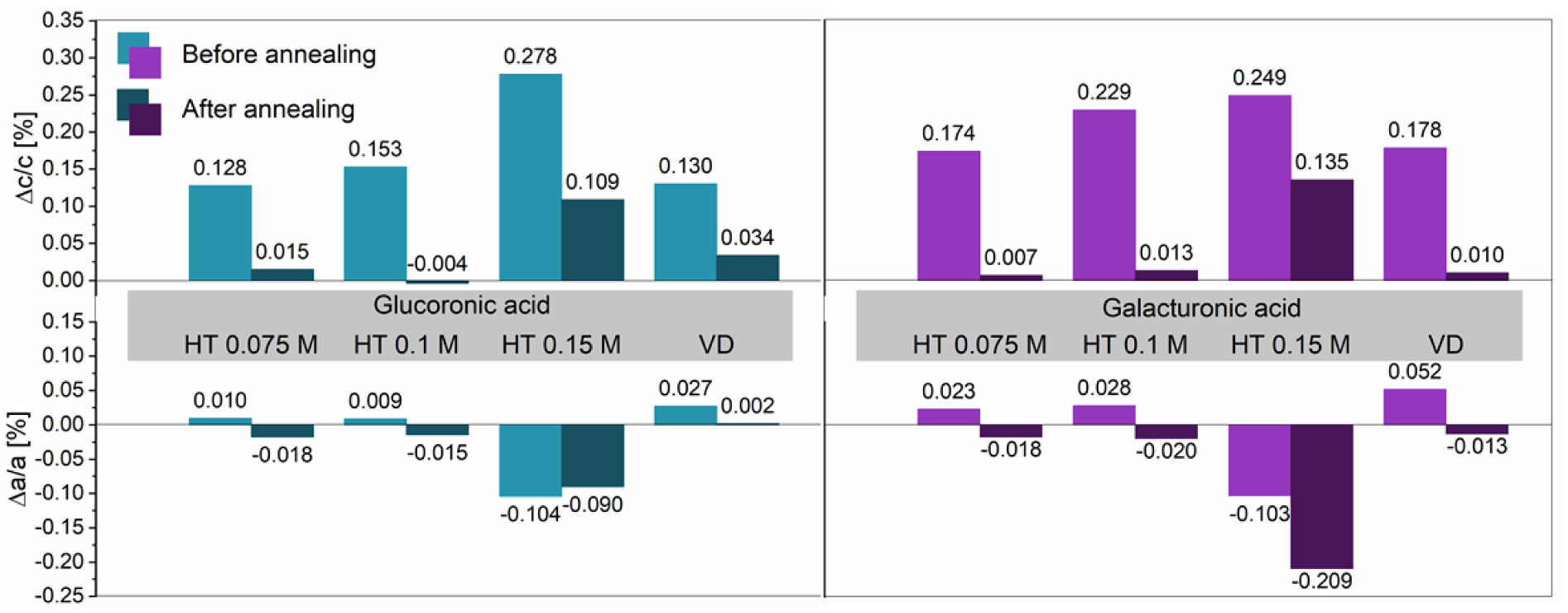
Lattice distortions induced in calcite samples grown via the HT or the VD method and in the presence of 0.075, 0.01 or 0.15 M of GlcA (left) or GalA (right). Top – distortions in *c*-parameter; bottom – distortions in *a*-parameter.

Broadening of diffraction peaks after annealing is a well-established phenomenon in biogenic crystals as well as when organic molecules become incorporated into the crystalline host.^[77]^ The main reason for this is that after annealing the organics decompose and produce new interfaces, which reduce the coherence size (crystallite size) and increase the microstrain fluctuations. To measure these microstructural changes we performed line-profiling analysis as previously described.^[77]^ Analysis in the case of the (104) diffraction profile indeed confirmed the reduction in crystallite size and increase in microstrain fluctuations that resulted from annealing of the samples containing incorporated MSs (Table S2).

To measure the concentrations of MSs within the calcite we performed Total Organic Carbon analysis (TOC). This method enables us to detect the amount of carbon found solely in an organic compound and hence to precisely quantify the amount of MSs incorporated into the calcite crystal lattice. Because of the scarcity of available calcite material where MS concentrations were highest, we performed TOC analysis for calcite crystals grown under HT conditions in the presence of 0.075 M GlcA and 0.075 M GalA. According to the results, the amounts of incorporated GalA as well as of GlcA were as high as 1 wt%. Interestingly, even though in the case of the lowest MS concentration of 0.075 M, where the crystal morphology and lattice distortions were the least affected by MSs, we still detected relatively high amount of incorporation (1 wt%) for both of the MSs used. Previous reports on biogenic and synthetic calcite crystals containing organic inclusions have clearly verified the concept of a direct correlation between the magnitude of induced lattice distortions and the amount of incorporated organics.^[21–23]^ We may therefore conclude that also in the present case, the MS concentrations within crystals displaying larger lattice distortions clearly exceed 1 wt%.

We further aimed to define the direction of growth of the calcite crystals containing incorporated MSs. Using high-resolution transmission electron microscopy (HTRTEM) we analyzed the sample grown in the presence of 0.15 M of GlcA (which displayed the most prominent elongation and whose lattice parameters were altered the most). Measurements were obtained on the lamella taken from the short axis of the crystal prepared by FIB (Figure 5, inset). The bright-field image (Figure 5a) reveals an undisrupted crystalline lattice. Electron diffraction thus confirmed the single-crystalline character of the crystal and its elongation along [211] (Figure 5b). In order to reassure the crystals are indeed single crystals, we performed single crystal diffraction (SCD) measurements utilizing a synchrotron source. Indeed, all three examined samples (calcite control, calcite-GlcA and calcite-GalA) revealed a single-crystalline characteristic. The measured lattice distortions, according the lattice parameters obtained via SCD, together with representative SCD frames, are presented in Figure S6.

**Figure 5.**
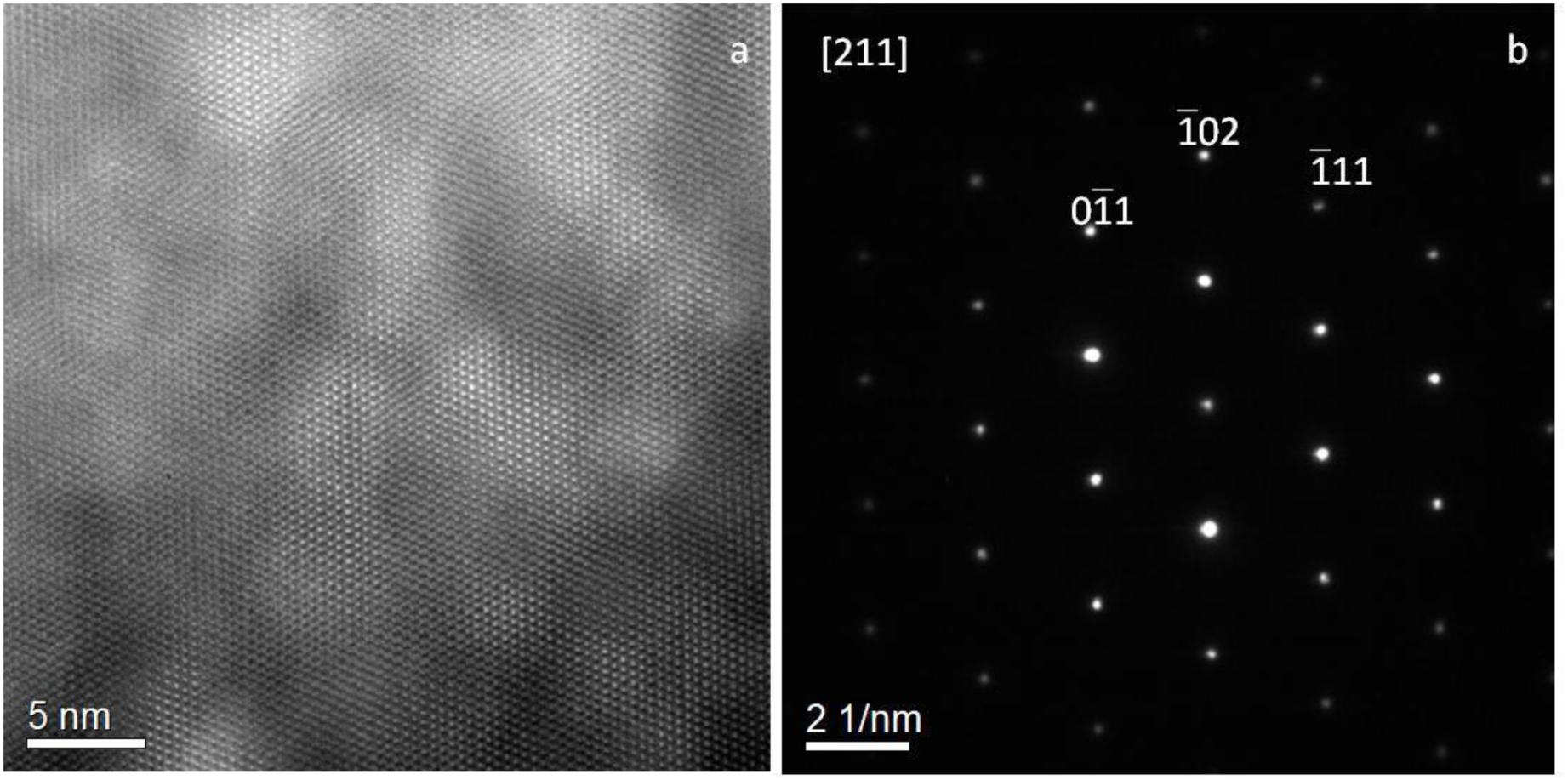
HRTEM analysis. (a) Bright field image of calcite with GlcA. (b) Electron diffraction represent the single-crystalline character of the crystal.

## CONCLUSIONS

To gain a better understanding of calcium carbonate biomineralization and to emphasize the key role played by saccharides moieties in this process, we investigated the incorporation of MS molecules into the crystalline structure of calcite. We studied the growth of calcite crystals under ambient and hydrothermal conditions. In the case of both growing methods we showed, for the first time, that incorporation of MSs into the calcite lattice indeed takes place, and that it affects both the morphology and the microstructure of the crystals. Importantly, we showed that as in the case of amino acid incorporation, the incorporated MSs induce anisotropic lattice distortions in calcite crystals, and that the level of lattice distortions is higher for crystals grown under HT than under ambient conditions. Moreover, we detected relatively high concentrations of occluded MSs, a finding never previously reported. Our results may have essential implications for assigning saccharides an unsuspected intimate interaction - at molecular level - with calcium carbonate crystals. We cannot exclude that such interactions may also contribute to stabilize transiently amorphous calcium carbonate phases. They may also have an important impact on the development of innovative composite materials. In the future, it will be interesting to check on fossil biominerals whether this interaction is stable over time, *i*.*e*., whether fossil biominerals demonstrate a relaxation of their lattice parameters (due to the saccharide degradation) or whether they still exhibit lattice distortions caused by preserved acidic saccharides.

## EXPERIMENTAL SECTION

### Synthesis

CaCO_3_ crystals were grown in the presence of three MSs: D-(+)-GlcA monohydrate (reagent grade, 97%), D-GalA (reagent grade, 98%) and D-Glucose (reagent grade, 98%), all purchased from Sigma-Aldrich. Deionized water was used for the preparation of all solutions.

#### - Vapor diffusion (VD) method

GalA (0.15 M) or D-(+)-GlcA (0.15 M) was added to an aqueous solution of CaCl_2_ (0.15 M, 100 mL) and stirred until the MS was dissolved. The resulting solutions in glass vails were placed in a desiccator together with a CO_2_ source (4 g of ammonium carbonate), and left for 48 h to allow synthesis to proceed. Control CaCO_3_ was prepared using the same synthesis procedure but without MSs.

#### - Hydrothermal (HT) method

D-Glucose 0.15 M or GalA or D-(+)-GlcA in various concentrations (0.075 M, 0.1 M and 0.15 M) was added to an aqueous solution of Na_2_CO_3_ (0.15 M, 100 mL each) and stirred until completely dissolved. The CaCl_2_ solution (0.15 M, 100 mL) and Na_2_CO_3_ solution containing dissolved MSs were mixed rapidly and then placed into an autoclave for 4 h at 134 °C and under water pressure of ∼0.3 MPa. As a control, CaCO_3_ was prepared according to the same synthesis procedure but without MSs. The finally obtained solutions were filtered through filter papers (2.5 µm) and the recovered powders were washed several times with deionized water.

### High-resolution powder X-ray diffraction (HRPXRD)

Powder diffraction experiments were conducted at the ID22 beam line of the European Synchrotron Research Facility (ESRF), Grenoble, France. The experiments were carried out at wavelengths of 0.39999(168) Å and 0.49602(95) Å. Each sample was loaded into a borosilicate capillary, and their X-ray diffraction patterns were collected before and after ex-situ annealing (300°C, 3 h in air).

### Focused Ion Beam (FIB) and High-Resolution Transmission Electron Microscopy (HRTEM)

A cross section of a calcite **sample containing** GlcA was prepared using a focused gallium ion beam (Helios NanoLab G3 UC) at a voltage of 30 kV and a current of 21 nA. Sample preparation was followed by thinning, done while maintaining a constant voltage and gradually decreasing the current from 0.43 nA to 80 pA. Polishing was done at 5 kV and 15 pA. The FEI Titan 80−300 FEG-S/TEM system was used in bright-field TEM mode on the FIB-cut lamella, as well as to perform electron diffractions.

### XRD data analysis

Lattice parameters of the CaCO_3_ crystals were extracted via the Rietveld refinement method by employing GSAS-II software^[78]^. Lattice distortions were determined with high precision as in our previous studies.^26^ Crystallites sizes and microstrain fluctuations were determined by fitting the (104) peak, the most intense peak of calcite (Figure S2), to a Voigt function.^[77]^

### Total organic carbon (TOC) analysis

Calcite crystals were bleached in 25% v/v sodium hypochlorite (NaOCl) for 5 min to remove surface-bound organics. This treatment preserves only the intra-crystalline organic molecules. The crystals were centrifuged and washed several times with deionized water. TOC values were determined at Aminolab, Ness Ziona, Israel. In a typical measurement, a bleached sample was dissolved in 0.2 M HCl, and phosphoric acid (H_3_PO_4_) was then added to completely remove the total inorganic carbon. <u>Oxidation</u> of organics to CO_2_ was initiated by adding peroxydisulfuric acid (H_2_S_2_O_8_). The amount of CO_2_, which is directly related to the amount of TOC in the sample, was detected by Fourier Transform Infrared Spectroscopy (FTIR).

### Single-crystal X-ray diffraction (SCD)

Diffraction data of single calcite crystals have been collected at the ID23-1 beamline of the ESRF. The collection was done at room temperature using a wavelength of 0.85 Å, an oscillation angle (Δφ) of 1° and a detector-sample distance of 129.86 mm. The detector was a Pilatus 6M. The data have been processed by the software XDS^[79]^ imposing P1 as space group. The unit cell axis and the data collection statistics are reported in Table S3.

## SUPPORTING INFORMATION

Supporting Information is available from the Wiley Online Library or from the author.

## ACKNOWLEDGEMENTS

This project was partially funded by the European Union Horizon 2020 Research and Innovation Program under the Marie Skłodowska-Curie Grant Agreement no. 642976-NanoHeal Project. We acknowledge the European Synchrotron Radiation Facility for provision of synchrotron radiation facilities at the beamline ID22.

## ABBREVIATIONS

FIB: focused ion beam
GalA: galacturonic acid
GlcA: glucuronic acid
HRPXRD: high-resolution powder X-ray diffraction
HRSEM: high-resolution scanning electron microscopy
HRTEM: high-resolution transmission electron microscopy
HT: hydrothermal (16)
MSs: monosaccharides
PSs: polysaccharides (11)
TOC: total organic carbon
VD: vapor diffusion.

## Notes

### Competing Interest Statement

The authors have declared no competing interest.

### Summary of Updates

references corrected

